# Targeting Colorectal Cancer Liver Metastasis Through Repurposing Metabolic and Immune Inhibitors : A Theoretical Study

**DOI:** 10.1101/2025.02.12.637833

**Authors:** Madhurima Mondal, Aditya Lahiri, Aniruddha Datta

## Abstract

Liver metastasis from colorectal cancer is a major cause of death in patients with advanced colorectal cancer (CRC) and remains a difficult challenge in oncology. Inspite of commnedable developments in medical inventions, the complexity and poor prognosis of metastatic stage, alliterative strategies are required to address Colorectal cancer liver metastasis (CRLM). This paper presents a computational study on drug repurposing for CRLM using a Boolean Network model. We comprehensively analyze CRLM signaling pathways such as WNT, PI3K/AKT/mTOR, MAPK, Hedgehog, TGF-*β*, NF-kB, NOTCH and HGF and target them with metabolic and immune inhibitors. This study utilizes a computational analysis to evaluate single and multi-agent treatments across scenarios involving single and multiple genetic mutations. Our findings highlights the efficacy of metabolic inhibitors such as Simvastatin, Metformin, and predict compatible partner drugs to enhance their effiacy. Additionaly, we predict possible improvements for CRLM treatment using an immunotherapy drug like Pembrolizumab. Overall, this paper provides new insights into drug repurposing to improve CRLM treatment outcomes.

## Introduction

Colorectal cancer (CRC) is the third most frequent kind of cancer globally and a common cause of cancer-related mortality [1]. The American Cancer Society projected 152,810 new CRC cases in the United States for 2024, comprising 46,220 cases of rectal cancer and 106,590 cases of colon cancer [2]. Even though CRC develops in the last two parts of the large intestine, i.e., colon and rectum, distant metastases are common [1]. The liver is the most common site for CRC metastases, with approximately 50% of CRC patients developing liver metastases (CRLM) during the course of their disease [3–5]. Resistance against first-line and second-line therapies is common among metastatic CRC patients. Despite the advancements in medical science, treating CRLM is yet an overwhelming task due to disease reoccurrence and dismal outcomes [6–9]. Thus, discovering new therapeutic strategies for CRLM is necessary. This study explores the optimal therapeutic agents and their combinations, focusing on immune and metabolic inhibitors as a strategic approach for repurposing existing drugs that enhance the care of CRLM patients.

Standard treatments of CRLM comprise surgeries, chemotherapy, stereostatic body radiation therapy, and adjuvant chemotherapy [10–12]. Even though surgical treatments are a benchmark for CRLM, only 20-30% of CRLM patients can go through surgeries constrained on tumor size, location, spread, and other medical conditions [10, 13, 14]. Systematic chemotherapy, intra-arterial chemotherapy, and local ablation therapies are suggested for patients with unresectable CRLM [14–16]. However, they do not offer the same survival benefits as surgical resection when these treatments are solely applied. According to available data, Stereotactic Body Radiation Therapy (SBRT) for CRLM has shown promising outcomes in terms of safety, local control (LC), and overall survival (OS) [11, 17, 18]. SBRT shows limitations on tumor size and location and can potentially harm adjacent tissues. Molecular targeted therapy is another key element in cancer treatments [19–21]. In order to create advanced treatment strategies, researchers identify possible genetic abnormalities and the resulting aberrant proteins that contribute to the development of particular cancers [22, 23]. Unlike chemotherapy, the targeted treatment medication affects the aberrant protein only. Standard targeted therapies include anti-angiogenic agents, epidermal growth factor receptor (EGFR) inhibitors, and immune checkpoint inhibitors [24–27]. Unlike several other cancers such as endometrial cancer, lung cancer, and gastric cancer, immunotherapy by itself did not show promising results in CRLM [28–31]. Alternative therapeutic options such as dietary supplementary are gaining traction in cancer treatment [32, 33]. However, nutritional supplements which offer a slow mechanism of action, may not be effective in treating aggressive metastatic cancers such as CRLM. Consequently, new drug discovery for CRLM still remains challenging, given the complexity of target-based drug development, the high cost, and the time commitment associated with developing new therapies. Identifying side effects of new drugs also requires years of preclinical and clinical trials. For diseases like CRLM, perioperative mortality can be up to 30%, and around 40% of patients experience recurrence within one year. Due these high risks, effective treatments are desperately needed and usually cannot wait for the classical timescale associated with medication discovery [34]. Because of these limitations, repurposing drugs with known safety profiles and new therapeutic indications, emerges as a necessary and affordable alternative for CRLM. When administered with current chemotherapeutics, anti-angiogenic drugs such as Bevacizumab have demonstrated promise in enhancing overall and progression-free survival in patients with CRLM [35]. While effective in treating hormone-sensitive tumors, including prostate and breast cancer, hormonal therapy can also reduce the risk of colorectal cancer. However, hormonal therapies do not offer much hope for CRLM as the metastatic initiation is less correlated to hormone-specific mechanisms [36].

Recently, a new direction of targeted therapy in metastasis has emerged with the drug repurposing through metabolic inhibitors [37–40]. Cellular metabolism is essential for responding to changes in a cell’s internal and external environment because it controls various molecular and biochemical activities [41]. To meet the increasing bioenergetic needs of overcoming complex hurdles throughout the cellular transit, cancer cells, in particular, modify their metabolic circuits during the epithelial-mesenchymal-transition (EMT). Thus, determining how metabolic inhibitors change the EMT and cancer signaling pathways, would provide important and susceptible treatment targets [38, 42, 43]. On the other hand, immunotherapy is also getting attention in cancer treatments such as endometrial, lung, and gastric cancers; however, it did not show excellent outcomes as stand alone treatment in CRLM [26, 29–31, 44]. Hence, a comprehensive understanding of cancer metastasis signaling pathways with metabolic and immunotherapy inhibitors represents an essential step towards designing effective combinatorial therapies.

Cancer, a collection of malfunctions such as uncontrolled proliferation in the cell cycle regulatory system, is driven by deleterious genetic and epigenetic mutations [45]. Hence, studying the molecular pathways within the cell where these mutations occur is insightful [33]. The recent breakthroughs in the development of new generations of cancer therapies, such as those enabled by next-generation sequencing (NGS) and multispectral analyses and understanding of cancer signaling pathways, have significantly improved the knowledge of the molecular mechanisms in cancer [32, 33, 46]. Researchers have discovered intricate cell network formation through several cancer signaling pathways, which provide the information needed to pinpoint the molecular targets for cancer treatment [47]. The last phase of neoplastic progression, namely metastasis, occurs as genetically unstable cancer cells adapt to a distinct tissue microenvironment, giving rise to new tumors in distant organs and facilitating the spread of cancer from its primary site [48–51]. Metastatic development happens through several advancements in cancer progression. The metastatic cascade is the term used to describe the intricate series of actions involved in this process. EMT tumor cell dissociation, local invasion, intravasation into blood and lymphatic vessels, transport through the circulatory system, extravasation at secondary sites, and the establishment of micrometastases and macrometastases are the subsequent steps in the process, which starts with the creation of a premetastatic niche at the target site [50–53]. In this paper, we conducted a comprehensive analysis of the CRLM cancer signaling pathways that affect the metastatic cascade, with the goal of designing effective interventions to target the metastatic spread of cancer.

Despite pathway information being marginal and limited in nature, studying signaling pathways, may offer useful therapeutic clues for tumor progression brought about by genetic mutations in them. In the case of cancer, especially for metastatic stages, the pathway information is usually very limited. Numerous strategies have emerged for identifying therapies for CRLM by leveraging an understanding of gene interactions within metastatic pathways. These strategies encompass using established pathways from public databases, network-based methodologies, and the de novo construction of cancer-specific pathways [54]. In this paper, we utilize genetic regulatory networks to model multivariate interactions, use these networks to identify probable breakdown points that cause metastasis, and even suggest prospective therapeutic approaches involving combined targeted therapy. We employ one network-based methodology, namely Boolean Networks (BNs), to analyze the signaling pathways associated with CRLM. We leverage metabolic and immuno-inhibitors to target colorectal cancer liver metastasis by analyzing the associated signaling pathways through a BN model. Our results focus on the followings :

- We computationally predict promising combination therapies involving metabolic inhibitors for CRLM.
- We computationally show improvement options in immunotherapy for CRLM. The paper is organized as follows. In Section 2 we provide a concise introduction to

Boolean network modeling. Section 3 discusses the pathways and gene interactions relevant to CRLM. Section 4 details the mechanisms and intervention points of various drugs and inhibitors. Section 5 consists of the results obtained from the Boolean network analysis, including predicted therapeutic efficacies. Finally, Section 6 concludes the paper by discussing key insights obtained from the simulations and outlines the direction for prospective future research in CRLM therapy development.

## Boolean Pathway Modeling

In this paper, we represent biological systems as an integral graph of binary interactions between different biological components in a cancer pathway inside a cell [32, 55]. We build upton the idea of sequential interactions between genes and cross-talks within signaling pathways implicated in CRLM. CRLM signaling pathways influence various metastasis steps such as proliferation, apoptosis, prognosis, EMT, and angiogenesis. In healthy cells, signaling pathways function in harmony factors that mitigate pathway disruptions maintaining internal stability and proper cell cycle [32, 56, 57]. Analysis of cancer pathways, made up of a collection of genes that direct a specific biological function, are popular methods [58]. As seen in the KEGG database [59], the edges in a signaling pathway represent signals from one gene to another, and the nodes in the pathway represent genes and their products. Researchers have done intensive studies to understand and analyze these gene-gene interactions in cancer-related signaling pathways. Probabilistic graphical models [60], Next-generation sequencing [61], Microarrays [62], RNA sequencing [63], RNA interference [64], Deep learning [65], Machine learning [66], Boolean Networks [32, 56] are a few examples of the tools and techniques employed to carry out such studies. When looking for predictive relationships between a potentially large number of in a transcriptome (about 20000) using data from a small group of patients (about 1000), overfitting is commonly encountered in deep learning and machine learning models [67]. This limitation occurs due to the curse of dimensionality [68]. Additionally, due to the shortage of publicly available large-scale gene expression and patient data on CRLM, exclusively data-driven models may not be suitable for predicting gene interactions in CRLM. Consequently, we propose BN modeling, a data-independent method for this paper [32, 56, 69]. We build a BN model based on currently known protein-protein interactions in cancer signaling pathways from the existing literature. The BN modeling explains and analyzes the interactions of the drugs with the signaling pathways.

### Modeling signaling pathways

The boolean network model is equivalent to a state transition diagram [70] consisting of directed acyclic graphs [56]. The graph nodes represent the genes in the signaling pathway, and the edges between the nodes represent the directed signals between the relevant pair of genes. We consider two states of the genes: healthy and mutated (faulty). Thus, the nodes always assume binary values, either 1 or 0. In boolean models, a “state” is defined as a binary vector formed from simultaneously considering all gene values. Thus, for a boolean network with *n* nodes, the cardinality of the state space is 2^*n*^ [71]. The interactions between the genes, represented by nodes in the Boolean network, are given as boolean functions. The inputs to the boolean function for each node are the relevant interacting genes (one or multiple) and the (single) output for the Boolean function is the gene value associated with that node. Thus, each boolean function represents a Multi-input single-output (MISO) system. The inputs in the BN are the proteins, such as growth factors or tumor suppressors in each signaling pathway. These proteins are either on (1) or off (0). The final output of the BN consists of the reporter genes, which are responsible for tumor cell proliferation, apoptosis, EMT, metastasis, and angiogenesis. Thus, the Boolean network, taken as whole, represents a multi-input multi-output (MIMO) system. We illustrate the boolean functions in a biological system with a toy example in Fig 1a, based on the interactions among the inputs. In this paper, the terms we refer to “gene” and “node” are interchangeably used. In a BN, the state vector is a binary one based on measured genes - expressed or unexpressed, i.e., high or low [71]. In Fig 1a, we consider a biological signaling pathway with genes *a* through *f*. Here, in Fig 1a, genes *a* and *b* act simultaneously to form a dimer which activates gene *c*. The gene *c* acts as an inhibitor of gene *e*. Gene *e* is also affected by the gene *d*, and finally, gene *e* activates the gene *f*. Based on these rules of regulatory interaction, we can arrive at the boolean equivalent model shown in Fig 1c. In this figure, since genes *a* and *b* form a dimer to act on node *c*, the boolean function for node *c* = *a* ∧ *b*. Since *c* acts as an inhibitor for *e*, we use a NOT operator on *c* and feed it to node *e*. Since gene *d* also activates gene *e*, we use an inhibition by *c* on *e* and an OR gate to add the activation by *d*. Thus, the boolean function for node *e* = (~ *c*) ∨ *d*. Since node *e* also activates node *f*, we use a buffer to go from e to f, i.e., *f* = *e*.

**Fig 1.**
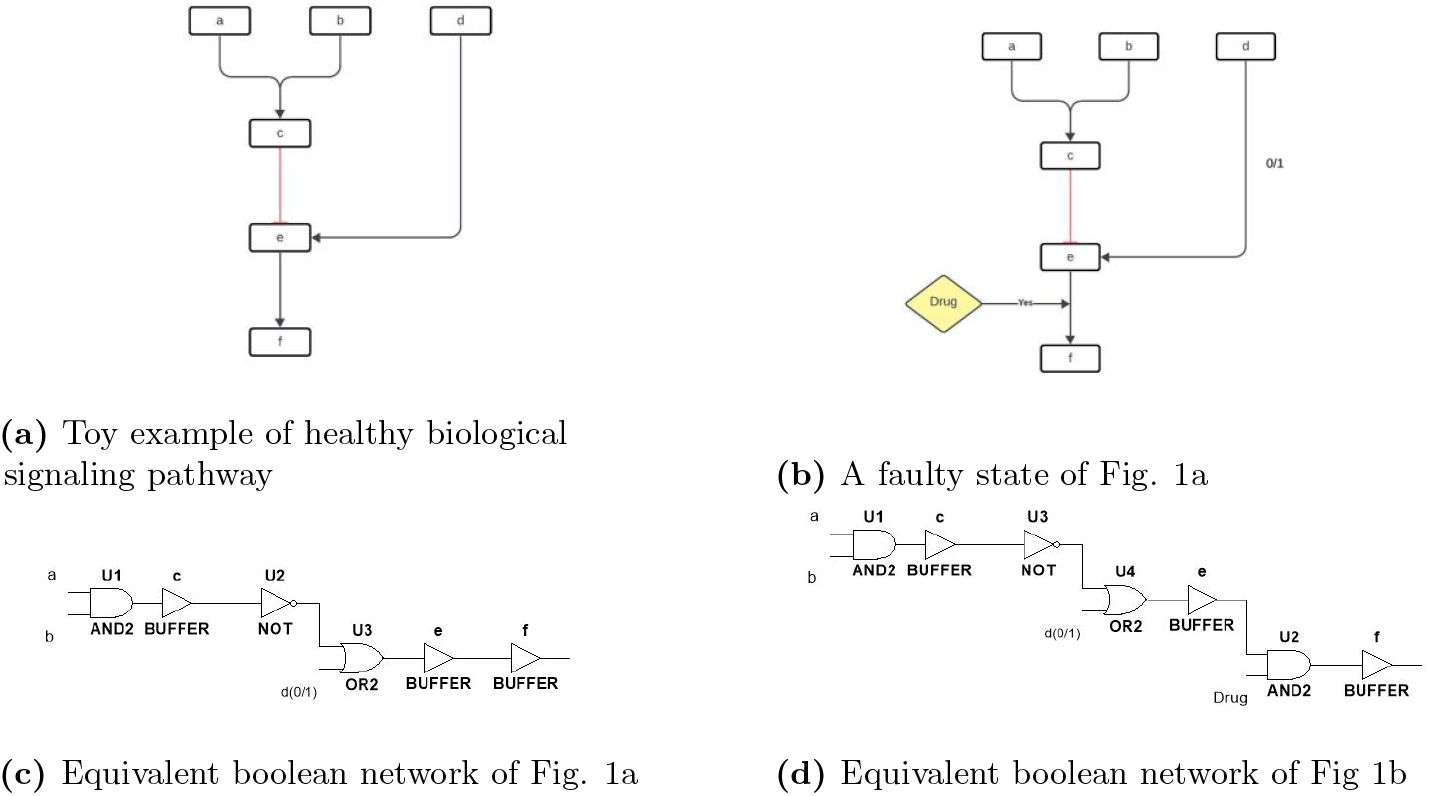
Toy example of healthy and faulty signaling pathways and their corresponding boolean network models

### Modeling abnormalities in signaling pathways

Genetic alterations in the signaling pathway can lead to cell abnormalities, and the development of tumors [32]. Therefore, studying the ablterations in our BN is of paramount importance here. A genetic alteration can happen due to hyperactive or mutated genes [32, 56]. We refer to these aberrant behavior caused by the mutated genes as “stuck at fault” state [56]. Growth factors or interacting genes cannot affect the expression level of a gene, once it is in a “stuck at fault” state. In other words, the state of a stuck-at gene is independent of the status of any other gene - 1 (active) or 0 (mutated). Fig 1b represents a genetically altered version of Fig 1a where gene *d* is mutated.

### Modeling corrective drug interventions in the signaling pathway

Targeted therapeutic agents bind to specific mutated genes according to their molecular properties. Unlike chemotherapy, targeted therapies are able to differentially affect cancer and healthy cells [32]. Targeted drugs bind to their target gene and consequently either suppress the gene or enhance the expression of the targeted gene. The “Drug” in Fig 1b represents a targeted therapeutic agent for restoring the normal behavior of the mutated gene *d*. We superimpose the effects of a drug on the BN of Fig 1b to obtain the BN of Fig 1d. The boolean functions from Fig 1d now include the effects of the Drug. This drug targets the gene *e*, opposes its inhibition, and activates health signals to *f*. Thus, the Boolean function of defining *f* changes to *f* = *e* ∧ (~ Drug).

## Signaling pathways in CRLM

Signaling pathways made up of proteins, genes, and transcription factors, control critical cellular processes like metabolism and differentiation through a coordinated network of molecular interactions. For a cell to function properly, it relies on extracellular biological signals through signal transducers [72]. This process starts with a ligand, such as hormones or growth factors, binding to a cell membrane receptor, which in turn, sets off a series of genetic signaling events through various proteins. In downstream signaling, this mechanism modulates transcription factors in the nucleus and alters the DNA expressions [73]. Disrupted expression of molecular components within intracellular signal transduction crucially impacts cellular functions, affecting normal growth, differentiation, neurotransmission, and the development of cells [74]. Researchers have investigated the existence of multiple alternative signaling pathways in cancer development and found the link between mutations in key driver genes and the metastasis of colorectal cancer [75]. Metastasis pathways often include additional elements to the primary tumor pathways depending on the metastatic site [76]. Building upon this foundation, this paper explores CRLM pathways to unveil the complex interplay of genetic signaling and pathway mutations that lead to the liver metastasis of CRC. Given the pivotal roles of such genetic mutations in the progression of CRC liver metastasis, we explore the mutations associated with common liver metastasis pathways in CRC, such as the WNT, TGF-*β*, PI3K/AKT/mTOR, MAPK, NF-kB, NOTCH, Hedgehog, and HGF-cMET pathways [77–84]. In this paper, our objective is to explore the complex interactions of these pathways under normal and mutated conditions and predict potential treatment strategies that can inhibit liver metastatic effects. A comprehensive graphical representation of relevant CRLM signalling pathways and their crosstalks are given in Fig 2, and a brief description of each of these pathways follows.

**Fig 2.**
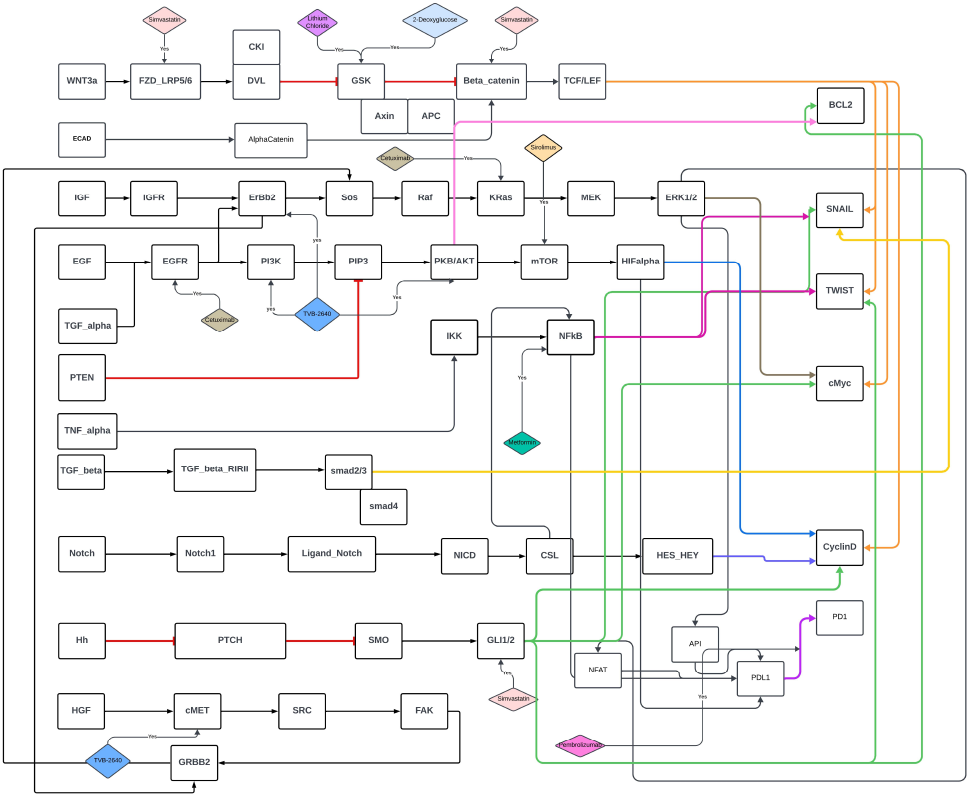
Relevant signaling pathways in CRLM

### WNT pathway

About 80%–90% of CRC patients have somatic mutations related to the Wingless-related integration site (WNT) pathway [77]. The main function of this pathway involves *β*-catenin, which controls the T-cell factor (TCF) and lymphoid enhancer-binding factor (LEF) in the nucleus. These are important for turning on genes that are responsible for cell growth, survival, and migrations [85]. About 70–80% of colorectal cancer cases have influence of this pathway with APC tumor-suppressor gene mutations. WNT signaling plays a vital role in both the initial oncogenic transformation in the colon and the subsequent metastatic development in the liver [77]. Overexpression of WNT3a in CRC stimulates the CRC progression through invasion and promotes EMT which leads to CRLM [86]. In the downstream, WNT pathway regulates genes such as CCND1, cMyc, SNAIL and TWIST. CCND1 and cMyc are responsible for the CRC progression and proliferation, whereas the SNAIL and TWIST are responsible for the EMT transition of the CRC cells [86, 87].

### PI3K/AKT/mTOR pathway

PI3K signaling mutations are frequently observed in liver metastatic or advanced stages of CRC patients [78]. Many in vitro and in vivo studies suggest that drugs that alter PI3K/AKT/mTOR pathway are important therapeutic targets [88]. Significantly lower PTEN expression, which normally suppresses PI3K, promotes liver metastasis in CRC patients [89]. Epidermal growth factor (EGF) [90], and transforming growth factor beta (TGF-*β*) initiate the binding with their respective growth factor receptors. These receptors, such as Epidermal growth factor receptor (EGFR) triggers the PI3K pathway in the CRLM [90, 91].The PI3K pathway affetcs reporter gene CCND1. that is responsible for driving prolifiration, and BCL2 which enhances tumor metastasis in CRLM [92].

### MAPK pathway

Many studies documented the role of the ERK/MAPK pathway mechanism in metastatic cancer [93]. This helps us to understand the effects of the ERK MAPK pathway on pathogenesis, progression, and metastasis of colorectal cancer [79]. The activation of the RAF/MEK/ERK signaling pathway is triggered by Insulin-like growth factor (IGF) and EGF [94]. Signals passed via protein kinase C (PKC), or RAS set off the ERK pathway signaling, leading to the activation of RAF, which starts a series of events activating MEK and subsequently, ERK [95]. Downstream effects of the RAS pathway include the activation of cMyc [79]. ERK also activates immuno-pathways affecting PDL1 and PD1 [84].

### TGF-*β* pathway

Transforming growth factor-beta (TGF-*β*) signaling is crucial in numerous physiological and pathological processes. Irregular TGF-*β* signaling pathway is key in advancing metastatic colorectal cancer (mCRC) [80]. The TGF-*β* signaling pathway is pivotal in a range of biological functions, like proliferation, differentiation, apoptosis, mobility, the epithelial to mesenchymal transition (EMT), changes to the extracellular matrix (ECM), blood vessel formation (angiogenesis), and immune responses within cells [80, 96]. TGF-*β* triggers the TGF-*β* receptor proteins, eventually activating SMAD2/3 [80]. This complex moves into the nucleus, triggering the activation of the EMT gene SNAIL [97].

### NF-kB pathway

Nuclear factor-kB (NF-kB) plays an important role in the progressions of CRC through proliferation, angiogenesis, and EMT [81]. Tumor necrosis factor-alpha (TNF-*α*) and other cytokines, which are involved in CRC liver metastases, activate catalytic subunits such as IKK-*α* and IKK-*β*, which further activate NF-kB [98, 99]. This pathway affects the EMT genes SNAIL and TWIST, which play a critical role in liver metastasis [100].

### NOTCH pathway

Abnormal Notch signaling, along with genetic mutations in Notch signaling-associated factors, impact the tumorigenicity and proliferation of cells in a variety of cancers, including CRC, where its inhibition can disrupt CRLM development [82, 84]. The human Notch system comprises several NOTCH receptors (Notch1); the Notch1 receptor releases NICD, and the expression of the ligand Jagged1 is upregulated. In the downstream, this pathway affects CSL and HES-HEY. These involve the transcriptomic genes, such as CCND1 and EMT gene TWIST, responsible for liver metastasis in CRC [84].

### Hedgehog pathway

The Hedgehog (Hh) signaling pathway plays a key role in proliferation, EMT and stell cell renewal in CRLM [83, 101]. The Hh signaling pathway begins with the production and secretion of Hh ligands [102]. Hh ligands bind and inhibit the transmembrane receptor PTCH which subsequently activates SMO [103]. SMO is responsible for affecting the transcription factor Gli activation [104]. Gli affects reporter genes CCND1, TWIST, SNAIL, and BCL2 [56, 83, 101].

### HGF-cMET pathway

HGF/c-Met signaling pathway promotes metastasis of cancer cells by regulating diverse downstream prometastatic effector molecules and enhancing prolifieration, migration and invasion [84]. Hepatocyte growth factor (HGF) receptor Met is positively correlated with the tumor stages of colorectal cancer liver metastases [105]. c-Met pathway affects reporter genes c-Myc and PDL1 through a crosstalk with the MAPK pathway [59].

### Drug’s intervention with CRLM signaling pathways

In this paper, we consider immunotherapy drugs and metabolic inhibitors for drug repurposing in CRLM. Each of these drugs, has it’s own target i.e. intervention node in different CRLM signaling pathways. Table 1 showcases the list of drugs, and summarizes targeted pathways, intervention nodes of the drugs, their standard use, and relevance towards CRLM signaling pathways.

**Table 1.** Drugs and Their Relation to Pathways in CRLM.

| Drug | Pathway;Target | FDA approval | Relevance to CRLM |
| --- | --- | --- | --- |
| Pembrolizumab | PD1; PDL1 [5] | FDA approved for immunotherapy in several cancers [136] | Inhibits immunosuppression, providing therapeutic benefit in CRLM. |
| Metformin | NF-kB; NF-kB [113] | FDA approved for type 2 diabetes [137] | Inhibits pro-inflammatory pathways, offering potential as a repurposed drug for CRLM. |
| Cetuximab | PI3K, MEK/ERK; EGFR, KRas [115, 118] | FDA approved for metastatic and recurrent head and neck cancer [138] | Reduces tumor growth in KRas wild-type CRLM by targeting EGFR signaling. |
| Simvastatin | WNT, Hedgehog; β-catenin, FZD, GLI1 [123–125] | FDA approved for dyslipidemia [38] | Inhibits multiple oncogenic pathways, suppressing tumor growth and metastasis. |
| TVB-2640 | PI3K, MEK/ERK, HGF ; PI3K, AKT, ERBB2, c-MET [129, 130] | FDA approved for noncirrhotic metabolic dysfunction-associated steatohepatitis [139] | Targets metabolic dependencies, reducing tumor growth and proliferation in CRLM. |
| Lithium Chloride | WNT;*GSK3β* [132] | FDA approved for manic illness [140] | Potentially disrupts tumor progression by modulating WNT regulatory pathways. |
| Sirolimus (Rapamycin) | PI3K;mTOR [133] | FDA approved for lymphangioleiomyomatosis (LAM) [141] | Modulates signaling pathways critical for CRLM tumor growth and progression. |
| 2-Deoxyglucose (2DG) | WNT;*GSK3β* [135] | - | Targets metabolic vulnerabilities, inhibiting tumor progression through WNT signaling. |

#### Pembrolizumab

Pembrolizumab is a FDA approved immunotherapy drug showing promising results in gastric, ovarian and endometrial cancer [106–108]. Pembrolizumab, an anti-programmed cell death (PD1) drug, affects PDL1 expression [5]. This immuno-inhibitor also affects several pathways such as MEK/ERK, NOTCH and thereby impedes tumor growth in colorectal cancer and provides therapeutic potential in CRLM [5, 109].

#### Metformin

Metformin, primarily known as an anti-diabetic drug reduces hepatic glucose metabolism and exerts anti-cancer effects by inhibiting the NF-kB signaling pathway [110–112]. Metformin inhibits invasion by targeting [113]. This inhibition reduces inflammation and tumor-promoting gene expression, making metformin a potential drug for repurposing in CRLM treatment [114].

#### Cetuximab

Cetuximab, another popular glycolysis inhibitor is an anti-EGFR monoclonal antibody that inhibits EGFR signaling by competitively binding to the receptor with higher affinity than TGF-*α* [115–117]. It inhibits EGFR and reduces tumor progression in KRas wild-type metastatic colorectal cancer, making it a key agent in CRLM therapy [115, 118].

#### Simvastatin

Another well known metabolic inhibitor for hypolipidomia is Simvastatin [119–121]. It targets multiple pathways, including Hedgehog and Wnt signaling. It reduces the expression of key proteins like *β*-catenin [122, 123]. Statins also affects the levels of FZD-LRP complex and targets GLI1/2 [124, 125] These effects contribute to the suppression of tumor growth and metastasis, making simvastatin a versatile therapeutic option for CRLM [126].

#### TVB-2640

TVB-2640 is a fatty acid synthase (FASN) inhibitor that disrupts cancer cell metabolism [127]. FASN inhibition reduces AKT phosphorylation levels and impairs ERBB2 expression, effectively limiting tumor growth [128, 129]. This FASN inhibitor targets ERBB2, PKB/AKT, PI3k and cMET [129, 130]. This mechanism highlights TVB-2640’s utility in addressing the metabolic vulnerabilities in CRLM [131].

#### Lithuium Chloride

Lithium chloride acts as an inhibitor for the enzyme glycogen synthase kinase-3 beta (*GSK*3*β*), thereby modulating key pathways involved in cancer progression [132]. By inhibiting *GSK*3*β*, lithium chloride can influence WNT signaling and other regulatory pathways, making it a potential therapeutic intervention for CRLM.

#### Sirolimus

Sirolimus, also known as rapamycin, is a potent mTOR inhibitor and a key regulator of cellular metabolism and growth [133]. Sirolimus is a metabolic inhibitor affecting lipid accumulation [134].

#### 2-Deoxyglucose

2-Deoxyglucose (2DG) is a glycolysis inhibitor that impacts WNT pathway activation by targeting *GSK*3*β* and forms a critical regulatory nexus within glycolysis [135].

### Simulation and Analysis

#### MATLAB Simulation

A BN representing the CRLM cell, operates as a multi-input multi-output (MIMO) digital circuit. A properly functioning BN system mimics the signaling pathways of a healthy cell. Healthy inputs direct the outputs in the healthy state. On the other hand, a malfunctioning BN represents the aberrant signaling of a malignant cell and generates outputs different from those of a healthy cell. In summary, the input-output behavior of a malignant cell BN differs from that of a healthy cell. This paper’s CRLM-BN network includes 11 inputs, 38 intermediate genes, and five outputs. For the healthy state of the BN, the tumor suppressors are active (state 1), while all other inputs are inactive (state 0). In our BN, the outputs act as cancer promoters, represented by state 0 in a healthy state. Fig 3 represents a healthy cell containing relevant CRLM pathways. Thus, in Fig 3, all inputs are healthy, and cancer-promoting outputs are turned off. One of the inputs, ECAD, is a key cell-cell adhesion protein that inhibits cell detachment and thus maintains tissue integrity. In signaling contexts, ECAD interacts with the WNT pathway by facilitating the activation of WNT signaling through its interaction with beta-catenin. However, to reflect the suppression of WNT pathway activation in a healthy state, ECAD is set to 0 in the boolean model.

**Fig 3.**
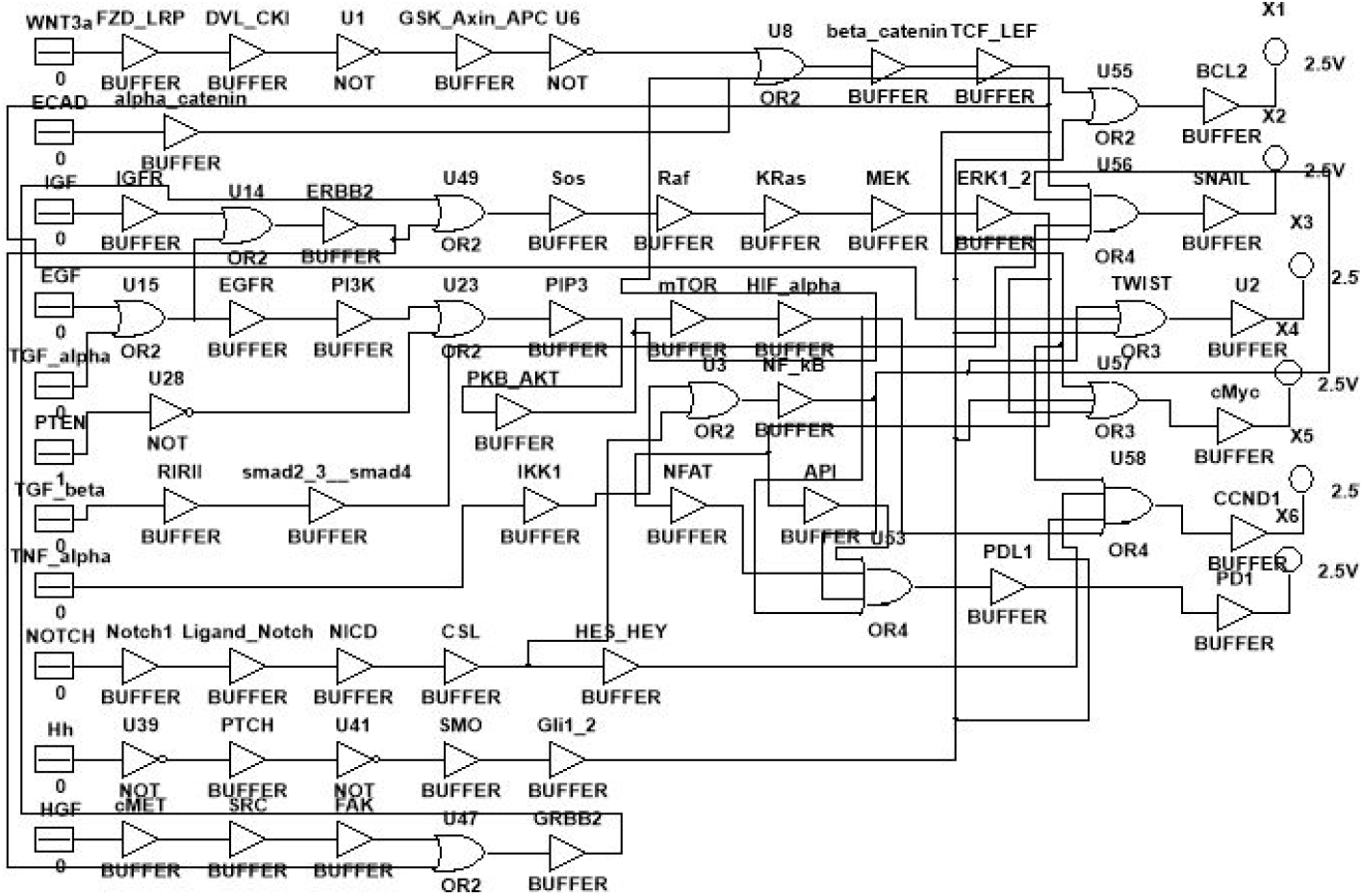
Boolean Network representation of CRLM pathways

Next, we apply different gene mutations inside the BN to generate a corresponding faulty output representing all possible malignant states that differ from the healthy state. At this stage, we apply the drugs in the BN to measure how each drug helps restore the healthy state of the malignant cells. We measure a drug’s efficacy by calculating the deviation of the drugged output state in a mutated cell from that of the healthy output state. We refer to this deviation as the size difference (SD) score. In general, the SD score between two binary vectors quantifies the dissimilarity between the vectors, and the higher score highlights a higher degree of dissimilarity. In this paper, we utilized a normalized mean SD (NMSD) score to consider the average over all possible SDs. Furthermore, the larger the NMSD, the larger the deviation from the healthy state, reflecting more extensive cellular abnormalities due to elevated proliferation, diminished apoptosis, and increased risk of cancer. Larger predicted NMSDs in the presence of by therapeutic agents that the particular therapeutic agent is less effective in restoring normal cellular processes and mitigating the effects of genetic mutations. On the other hand, a small NMSD indicates that the drugged state output is closer to the healthy state output, which reflects the therapeutic agent’s efficiency in mitigating mutations, reducing cancerous risks, and achieving cellular balance, thereby quantifying its effectiveness research in assessing cellular health.

To define NMSD mathematically, consider two binary vectors of dimension *n* i.e. **v**_1_ = (*v*_11_, *v*_12_, …, *v*_1*n*_) and **v**_2_ = (*v*_21_, *v*_22_, …, *v*_2*n*_). A confusion matrix captures the relationship between the elements of these vectors as follows:

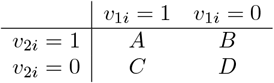

In the confusion matrix, the entries are defined as follows:

- *A*: The count of cases where the *i*-th elements of both vectors **v**_1_ and **v**_2_ are 1 (*v*_1*i*_ = 1 and *v*_2*i*_ = 1).
- *B*: The count of cases where the *i*-th element of **v**_1_ is 0 and the *i*-th element of **v**_2_ is 1 (*v*_1*i*_ = 0 and *v*_2*i*_ = 1).
- *C*: The count of cases where the *i*-th element of **v**_1_ is 1 and the *i*-th element of **v**_2_ is 0 (*v*_1*i*_ = 1 and *v*_2*i*_ = 0).
- *D*: The count of cases where the *i*-th elements of both vectors **v**_1_ and **v**_2_ are 0 (*v*_1*i*_ = 0 and *v*_2*i*_ = 0).

After measuring the mean SD (MSD) scored for given fault (either one, two, or three) scenarios and drug combinations, we summed these values and normalized them by dividing by the maximum score observed. This ensures the consistency with original normalized MSD (NMSD) score range of 0 to 1. Thus,

- *A* and *D* represent the counts of matches between **v**_1_ and **v**_2_,
- *B* and *C* represent the counts of mismatches between **v**_1_ and **v**_2_.

Using the confusion matrix, we define the mean SD by dividing the SD i.e., total mismatches in the confusion matrix mentioned before by total number of elements in the matrix itself. The definition is as follows:

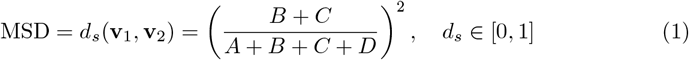

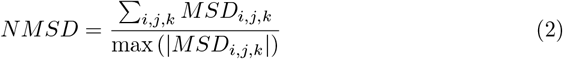

where *MSD*_*i,j,k*_ represents the SD score for specific fault dimension, (*i, j, k*). Normalized SD ranges between 0 and 1, where the value 0 corresponds to when the outputs are from a completely healthy state, mimicking a fault-free network. A normalized SD value of 1 indicates an untreated case or maximal deviation wherein the network fails to maintain normal cellular function.

In this paper, we implement the BN model using MultiSim to investigate the impact of various combinations of therapeutic agents on faulty genes (mutated genes) associated with CRLM cells. We simulated the network using MATLAB under 38 distinct genetic faults (mutations) and considered unhealthy states with combinations of one, two, and three faults alongside corresponding combinations of one, two, and three therapeutic agents. In essence, we computationally evaluate the efficacy of individual and combinatorial therapeutic strategies in the presence of single and multiple mutations. Integrating fault combinations and therapeutic agents allows us to identify effective therapeutic strategies for restoring healthy cellular functionality. Thus, we simulate the model to predict optimal therapies for 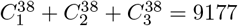 combination of faults, thereby providing robust results.

### Analysis

Our analysis highlights the varying performance of therapeutic agents and their combinations under different fault scenarios. We analyzed the results based on single-agent performance and their improvements in different combination therapies.

#### Therapy using a single drug

In this paper, the single-agent therapy scenario refers to the application of a single drug targeting particular genetic alteration within the signaling pathways. For this scenario, we also looked at one, two, and three fault situations. Excessive mutations can cause cellular plasticity, which allows tumor cells to adjust to therapeutic pressures. This makes it difficult to determine the true impact of individual or combination treatments. Therefore, we limited our analysis up to a maximum of three faults. Based on this setup, we found that in the case of single-agent therapy, the most effective drug, i.e., the drug with the lowest NMSD, is Simvastatin (targets FZD, *β*-catenin, GLI1/2) for all types of fault combinations. Fig 4 illustrates the efficacy of each drug under different fault combinations. The lower NMSD of Simvastatin demonstrates its exceptional ability to restore reporter genes closer to health state, regardless of the number of genetic faults present. However, the rise in NMSD of Simvastatin, with the increase in fault combinations, indicates a gradual reduction in its efficacy for cells with multiple mutations. This trend suggests that while Simvastatin performs well as a single drug, its therapeutic potential diminishes under higher mutational burdens.

**Fig 4.**
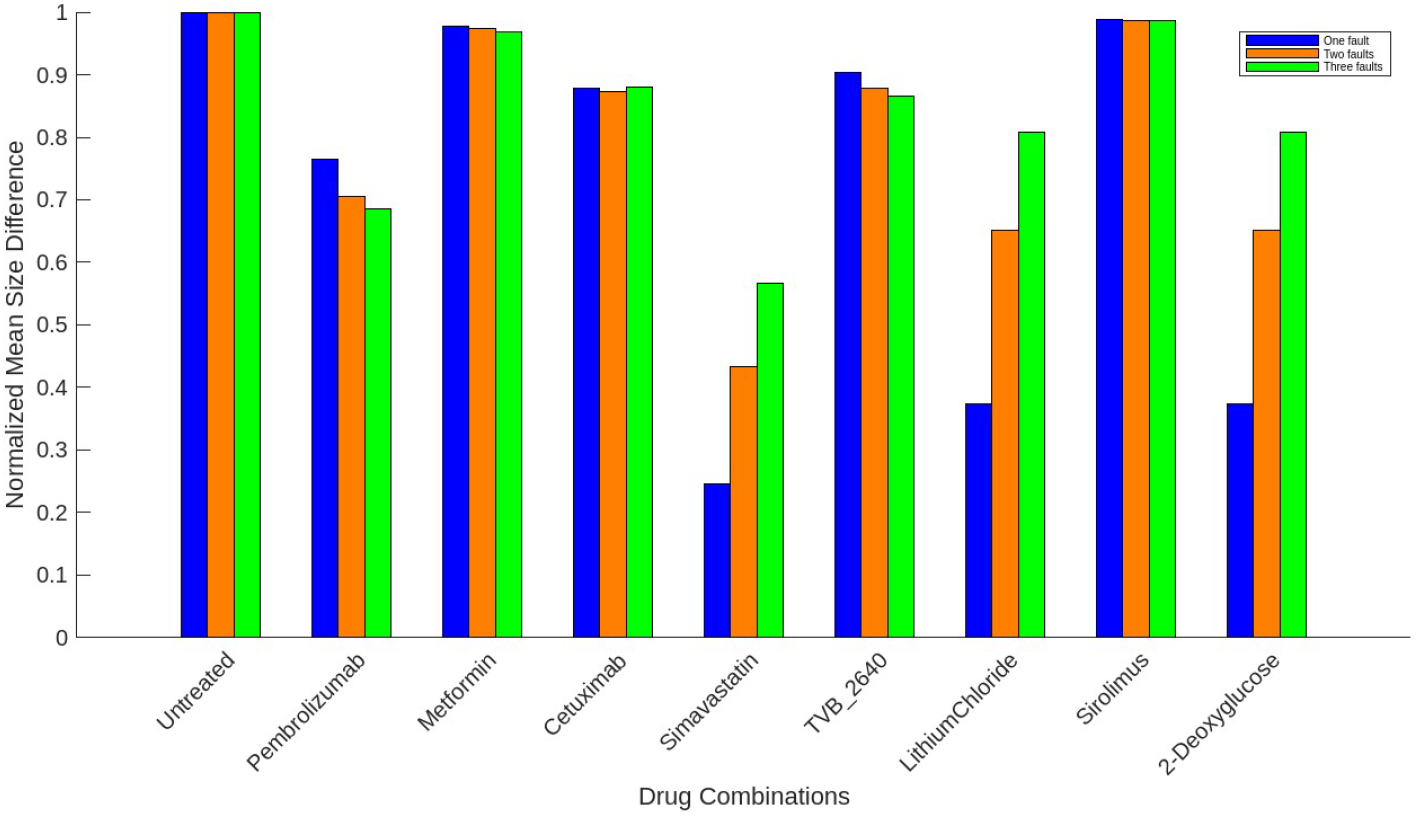
Normalized Mean Size Difference (NMSD) scores of drugs in single agent therapy in CRLM over one, two, and three faults.

Interestingly, Pembrolizumab, Cetuximab, TVB-2640, and Sirolimus perform better in two-fault combinations than single faults in signle agent therapy. This behavior could be caused by lethal pair (which refers to two genetic mutations that, when occurring together, result in more effective than targeting each mutation separately) effects among specific pairs of faults within the BN model [142]. Surprisingly, we observed no significant change in NMSD of Pembrolizumab from two-fault to three-fault combination scenario. This finding suggests that Pembrolizumab may reach its therapeutic saturation point by addressing two or more faults when used as a standalone agent. Whether this plateau holds for scenarios involving more than three faults in CRLM requires further investigation. Additionally, Cetuximab, TVB-2640, and Sirolimus show a slight increase in NMSD when transitioning from two-fault to three-fault scenarios. This observation suggests a reduction in the lethal pair effects at higher fault counts, possibly due to diminishing interactions among fault combinations.

On the other hand, Metformin exhibits consistently good performance across all three fault scenarios, ranking just below Simvastatin and Pembrolizumab in efficacy. However, the noticeable increase in NMSD for Metformin with higher fault combinations indicates its reduced efficacy in addressing cancers with multiple simultaneous mutations. Lithium Chloride and 2-Deoxyglucose showed similar standalone performance because of their same target points in the BN. We summarize these findings in Table 2, clearly comparing the NMSDs across the simulated drugs.

**Table 2.** Normalized Mean Size Difference (NMSD) scores of drugs in single agent therapy in CRLM over different fault combinations. Simvastatin shows the lowest overall NMSD, indicating the highest efficacy. The drugs are placed from lower to higher NMSD for one fault.

| Drug used | NMSD<br>for one fault | NMSD<br>for two faults | NMSD<br>for three faults |
| --- | --- | --- | --- |
| Untreated | 1.0000 | 1.0000 | 1.0000 |
| Simvastatin | 0.5201 | 0.5602 | 0.6117 |
| Pembrolizumab | 0.7368 | 0.6558 | 0.6591 |
| Metformin | 0.7585 | 0.8006 | 0.8371 |
| Cetuximab | 0.8607 | 0.8112 | 0.8423 |
| 2-Deoxyglucose | 0.8514 | 0.8715 | 0.9040 |
| LithiumChloride | 0.8514 | 0.8715 | 0.9040 |
| Sirolimus | 0.9876 | 0.9167 | 0.9455 |
| TVB-2640 | 0.9628 | 0.8594 | 0.8691 |

#### Therapy using drug multi-combinations

We define a multi-drug combination therapy scenario as the simultaneous application of more than one drug, each targeting specific genetic mutations inside the signaling pathways. For this scenario, we also considered one, two, and three fault situations to assess potential synergistic effects. Our BN model indicates that the simultaneous administration of two drugs often yields lower or comparable outcomes than when using the drugs individually for the respective fault combinations. Lowered NMSDs reflect improved efficacy in restoring reporter genes to their healthy state. Internal crosstalk of signaling pathways and the targeted intervention genes influenced by the combined drug actions contribute to these improvements. For instance, Fig 5d shows that Simvstatin with Pembrolizumab and with Metformin emerges as a notable combination for one, two, and three faults combinations. Though Simvastatin and Pembrolizumab show different trends in NMSD for higher faults when used as standalone agents, when administered together, the NMSD increases as the number of faults increases. This indicates that the combination’s efficacy reduces for cancers with multiple mutations as opposed to ones with only a single mutation. From Fig 5d Simvastatin did not improve when combined with Lithium Chloride and 2-Deoxyglucose. Lithium Chloride and 2-Deoxyglucose bind to the GSK-Axin complex in the WNT pathway, whereas Simavastatin binds to beta-catenin, a gene in the direct downstream of the GSK-Axin complex. This is why the combination of Lithium Chloride and 2-deoxyglucose with Simvastatin does not show synergistic effect. In the three drug scenario, Simvastatin showed a very good performance when administered along with Pembrolizumab and Metformin. Fig 6d shows the deatiled NMSD scores of three drug combinations that include Simvastatin.

**Fig 5.**
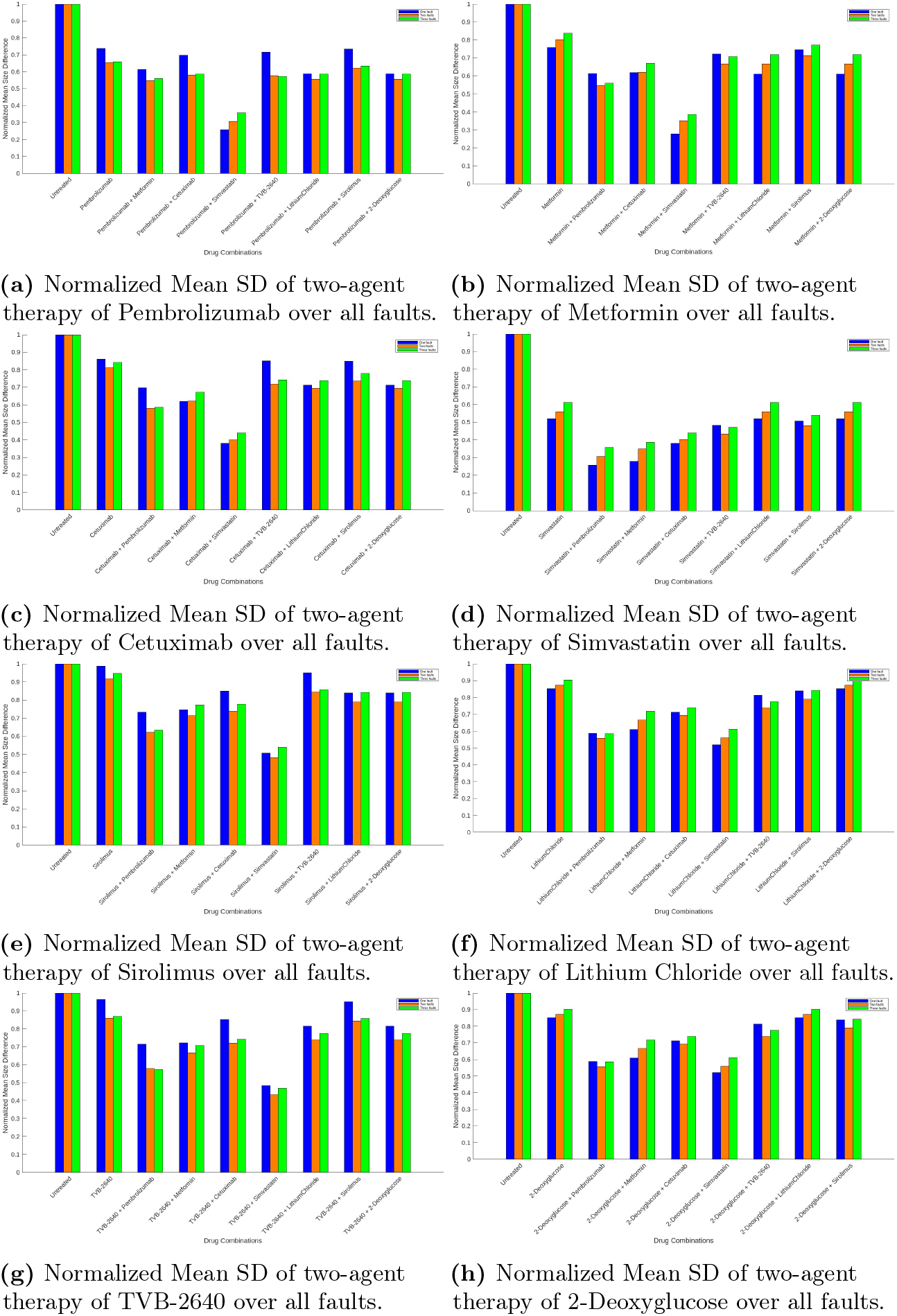
Normalized Mean Size difference scores of all two-agent therapies of each drug over one, two, and three fault combinations.

**Fig 6.**
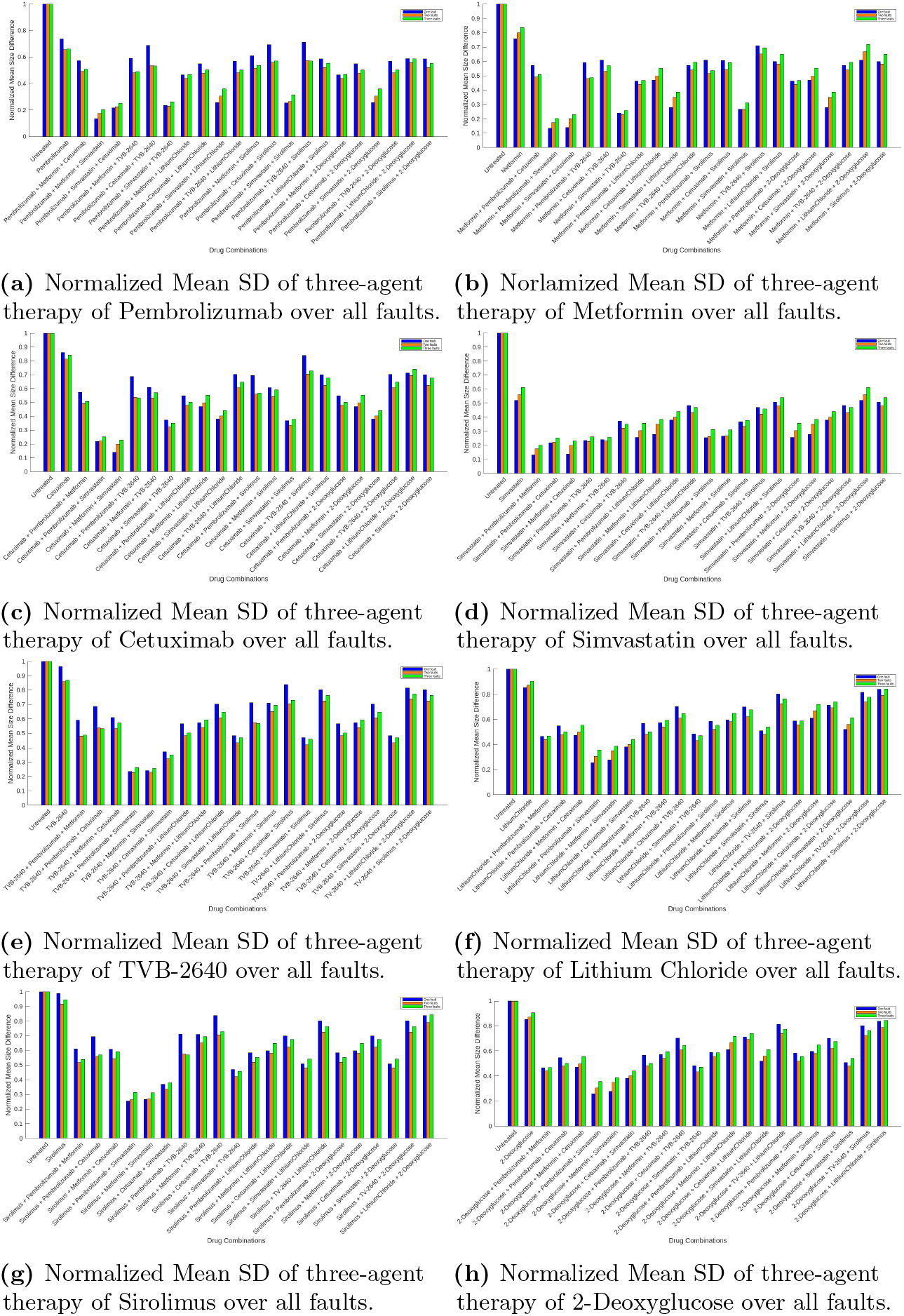
Normalized Mean Size Difference scores of all possible three-agent therapies of each drug over one, two, and three fault combinations.

Immunotherapy drugs like Pembrolizumab generally show limited success for CRLM due to the inherent resistance of colorectal cancer cells to immune modulation [143]. However, our computational simulations reveal that Pembrolizumab’s performance significantly improves when combined with metabolic inhibitors. For example, the combination of Pembrolizumab and Simavastatin, a metabolic regulator showcases the powerful synergy between immune checkpoint blockade and metabolic disruption. Pembrolizumab’s performances with other drugs in two-drug combination scenario are given in Fig 5a. Besides, the three-drug combination of Pembrolizumab with Simvastatin and Metformin yields exceptional results, as can be seen from Fig 6a. This highlights the possibility of using Pembrolizumab in combination with drugs that act on complementary pathways, like metabolism and cholesterol regulation, against CRLM.These findings emphasize the promise of combining immunotherapy with metabolic inhibitors to achieve superior outcomes in challenging cancer types like CRLM. We summarize the performance improvements of Pembrolizumab with metabolic inhibitors for single fault scenario in Table 3.

**Table 3.** Highest improvements in Pembrolizumab in combination therapy in the single fault scenario.

| Drug Combination | NMSD for One Fault | Percentage Improvement |
| --- | --- | --- |
| Pembrolizumab | 0.7368 | - |
| Pembrolizumab + Simvastatin | 0.2570 | 65.12% |
| Pembrolizumab + Metformin + Simvastatin | 0.1331 | 81.94% |

We then investigated the effectiveness of targeted treatments like Cetuximab. Cetuximab shows moderate standalone efficacy but performs better when combined with Simavastation for one, two, and three fault combinations. It achieves quite reasonable performance improvement when combined with the immunotherapy drug Pembrolizumab. However, pairing it with metabolic inhibitor drugs like Sirolimus and TVB-2640 shows limited synergy, with only marginal improvements. Interestingly, Cetuximab in combination with Pembrolizumab and Simvastatin performs well as a three-drug combination for one, two, and three fault combinations. This suggests a good therapeutic approach for complementary action among these drugs in targeting immune and signaling pathways. We summarize the NMSDs of two and three-drug combinations of Cetuximab in Fig 5c and Fig 6c respectively.

TVB-2640, which performs moderately as a single agent, shows excellent improvement while paired with Simvastatin for one, two, and three fault combinations. This pair predicts a lower NMSD within two-fault combinations, reflecting its increased efficacy. However, a slight increase in its NMSD when targeting triple-mutated cells, suggests reduced synergy effects, possibly due to the interaction of multiple mutations. Fig 5e showcases NMSDs of all possible two-drug combinations of TVB-2640 over one, two, and three faults. For three-drug combinations, TVB-2640 combined with Simvastation shows best results with Pembrolizumab and Metformin. TVB-2640 shows shows moderate synergistic effects with combinations of Sirolimus, Lithium Chloride and 2-Deoxyglucose. We summarize the three-drug combination performance of TVB-2640 in Fig 6e.

Fig 5b depicts that Metformin shows significantly improved efficacy when combined with Simvastatin. In the case of three drug combination, metformin performs best when combined with Simavastatin and Pembrolizumab. The combination of Metformin, Simavastatin and Cetuximab also achieves competitive performance. All possible three drug combinations are summarized in Fig 6b. Sirolimus,which performs average as a standalone drug, achieves better performance when combined with Simavastatin. Conversely, the combination of Sirolimus and TVB-2640 performs poorly. Fig 5g shows the NMSDs of all two drug combinations of Sirolimus. Fig 6d shows that the three-drug combination of Sirolimus with Simvastatin and Pembrolizumab achieves the best results combining immune checkpoint modulation and mTOR inhibition.

Our results show similar performances of Lithium Chloride and 2-deoxyglucose respectively in multi-drug therapy too. They bind to the same point of intervention in the CRLM pathway i.e. GSK, and hence show similar performances with other drugs as well. In our model, both of them showed limited efficacy when used in a standalone fashion, and even when combined with other drugs. Interestingly, in the case of one fault, they showed good performance when combined with Pembrolizumab. For three agent combinations, they showed the lowest NMSD when combined with Simvastatin and Pembrolizumab making the combination very attractive. Fig 5f and Fig 6f summarize the performances of Lithium Chloride for two and three-drug combinations respectively. Similarly, Fig 5h and Fig 6h summarize the NMSDs of 2-Deoxyglucose for two and three-drug combinations.

## Conclusion

In this paper, we presented a comprehensive analysis of optimal therapeutic agents for one, two, and three fault combinations in the context of CRLM using Boolean Network modeling. The study highlights critical insights into the synergistic effects of different metabolic inhibitor drugs and prospective improvements in immunotherapy in CRLM for cells with single and multiple mutations. As a standalone drug, Simvastatin emerged as the most effective therapeutic agent for one-fault, two-fault, and three-fault scenarios. Due to its pleiotropic effects, targeting multiple pathways, Simvastatin is a potent option for restoring normal cellular functionality in CRLM. Simvastatin showed a notable increase in efficacy when simultaneously administered with other medications, such as Pembrolizumab and Metformin. Hence, it makes a potent combination of targeted therapy. While, though immunotherapy showed excellent results in gastric and endometrial cancer, in our study it did not emerge as a reliable treatment option for CRLM thus supporting the existing literature on the efficacy of immunotherapy in CRLM [26, 29–31, 44]. Our study revealed an intriguing improvement of Pembrolizumab’s effectiveness when combined with metabolic inhibitors such as Simvastatin. The efficacy improvement of Pembrolizumab in combination therapy predicts a good synergy between immune modulation and metabolic intervention that can effectively counteract the immune evasion mechanisms of CRLM. Thus, this paper offers a promising avenue for overcoming CRLM’s resistance by repurposing metabolic and immune inhibitors, and can be a potential avenue of investigation for clinicians. Other metabolic inhibitors such as Cetuximab, Sirolimus, and TVB-2640 were demonstrated as potential repurposed drugs in CRLM depending on the fault combinations when used in the right combination. Cetuximab, targeting the EGFR pathway, reflected the potential for combination therapy by achieving improved performance with immuno-checkpoint inhibitors such as Pembrolizumab. Sirolimus, an inhibitor of mTOR, performed well with metabolic and cholesterol-regulating drugs. This finding takes advantage of complementary mechanisms that achieve better therapeutic results. TVB-2640 showed moderate performance, suggesting utility in specific synergistic contexts but underlining the importance of carefully selecting partner drugs. Metformin, a widely known metabolic regulator, has shown promising efficacy in CRLM, particularly when combined with immune modulators or cholesterol-regulating drugs.

Our BN model also captures the phenomenon of lethal pair effects, as observed with drugs like Pembrolizumab, TVB-2640, and Sirolimus. These drugs show higher efficacy in the presence of two mutations compared to just one, with their effectiveness remaining similar or slightly reduced when three faults are present. This trend may be attributed to the fact that these drugs target components downstream within their respective signaling pathways. Previous research such as [144] also witnessed such behavior among drugs.

The study found key limitations in drug combinations such as Lithium Chloride and 2-Deoxyglucose do not show synergistic effects in our pathway modeling. We can conclude that drugs that target the same mutations may not be useful in combination with targeted therapy. However, their comparable efficacy offers a valuable opportunity for precision medicine. It enables physicians to choose either 2-deoxyglucose or Lithium Chloride to address specific mutations in CRLM based on the patients’ drug resistance profiles, or side effects. This approach gives valuable input in drug selection for precision medicine.

This paper theoretically suggests, through a simplified Boolean Network model, an optimal treatment strategy for CRLM using metabolic and immune inhibitors. Though the paper concentrates on a few generic CRLM pathways, incorporating more specific pathways from patient data into our BN model could enhance prediction accuracy for precise therapeutic recommendations. Additionally, we did not account for any possible adverse effects of drug combinations. Therefore, addressing these shortcomings, including toxicity, resistance, and patients’ genetic profiles in modeling optimal therapy are potential areas of future research. Our work suggests potential laboratory experiments and drug discovery efforts that could enhance the benefits of combining immunotherapy and metabolic inhibitors for CRLM treatment. Such complementary beneficial drugs could bring about advancements in CRLM treatments. Also, investigating molecular properties of drugs that target lethal pair genes or downstream mutations in tumor cells with multiple mutations could be a topic for potential future work.

